# A nonhomologous end-joining mutant for *Neurospora sitophila* research

**DOI:** 10.1101/761130

**Authors:** Nicholas A. Rhoades, Elise K. Webber, Thomas M. Hammond

## Abstract

Disruption of the nonhomologous end-joining (NHEJ) pathway has been shown to increase the efficiency of transgene integration into targeted genomic locations of *Neurospora crassa* and other fungi. Here, we report that a similar phenomenon occurs in a second Neurospora species: *N. sitophila*. Specifically, we show that deletion of *N. sitophila mus-51* increases the efficiency of targeted-transgene integration, presumably by disrupting NHEJ. Researchers interested in obtaining the *N. sitophila mus-51*^Δ^ strains described in this study can obtain them from the Fungal Genetics Stock Center (FGSC).

## Introduction

The eukaryotic DNA repair machinery can prove to be a significant obstacle to fungal genome editing efforts. For example, while working with the model organism *N. crassa*, Ninomiya *et al.* (2004) found that targeted transgene integration (TTI)-success rates can be increased nearly fivefold (to 100%) by using a transformation host lacking a functional copy of the *mus-51* gene. MUS-51 is a critical NHEJ protein and the increased TTI-success rate caused by *mus-51* deletion is likely due to the inability of a MUS-51-deficient strain to preform NHEJ (for review, please see Chang *et al.* [2017]).

Several NHEJ-mutants have been constructed for use in *N. crassa* research (Ninomiya *et al.* 2004; Colot *et al.* 2006; Smith *et al.* 2016) and are readily available from the FGSC (McCluskey *et al.* 2010). While useful for researchers interested in modifying the *N. crassa* genome, the genus *Neurospora* contains many species (Nygren *et al.* 2011), some with unique genetic elements and/or biological processes, and the study of these other species could be aided by the availability of species-specific NHEJ mutants. For example, we are interested in *Spore killer-1* (*Sk-1*), a meiotic drive element found in natural populations of *N. sitophila* (Turner and Perkins 1979), and a readily available *N. sitophila* NHEJ mutant could make studies of *Sk-1* more feasible. Therefore, to aid our work on *Sk-1*, and to help others who wish to perform molecular genetic research on *N. sitophila*, we have constructed two *N. sitophila mus-51*^Δ^ strains. In the report below, we describe how the strains were constructed and we show that deletion of *mus-51* increases TTI-efficiency in *N. sitophila* as it does in *N. crassa*.

## Materials and Methods

### Strains and growth media

The key strains used in this study are listed in Table 1. Vegetative cultures were maintained on Vogel’s Medium N (VM) (Vogel 1956) with 2% sucrose and 1.5% agar unless otherwise indicated. Other media used in this study are BDS (VM with 2.0% L-sorbose, 0.05% D-fructose, 0.05% D-glucose, and 1.5% agar) (Brockman and De Serres 1963), SC-IAA (synthetic crossing medium with iodoacetate, as described by Ebbole and Sachs [1990]), TA (VM with 2.0% L-sorbose, 0.05% D-fructose, 0.05% D-glucose, 0.02% myo-inositol, 1 M sorbitol, and 1.0% agar); BA (VM with 2.0% L-sorbose, 0.05% D-fructose, 0.05% D-glucose, 0.02% myo-inositol, and 1.5% agar); and RM (VM with 2% sucrose and 1 M sorbitol).

**Table 1.** Strains used in this study

| Strain Name | Alternate names | Species / genotype |
| --- | --- | --- |
| FGSC 4746 |  | *N. sitophila* A (Tahiti) |
| W1426 |  | *N. sitophila* A (Italy) |
| FGSC ##### | ISU-4636 / HNR101.14.11 | *N. sitophila mus-51<sup>Δ</sup>::nat* A |
| FGSC ##### | ISU-4637 / HNR126.14.1 | *N. sitophila mus-51<sup>Δ</sup>::nat* A |
| P8-65 |  | *N. crassa csr-1<sup>Δ</sup>::hph* a |

### Vector construction

Vector 166 (v166) was designed to replace *mus-51* in FGSC 4746 and W1426 with a nourseothricin-resistance cassette (*nat*). The vector was constructed by Double-Joint (DJ)-PCR (Yu *et al.* 2004) using primers listed in Table 2. DJ-PCR was also used to construct vector 238 (v238), which was designed to replace *csr-1* in FGSC 4746 and ISU-4636 with a hygromycin-resistance cassette (*hph*).

**Table 2.**
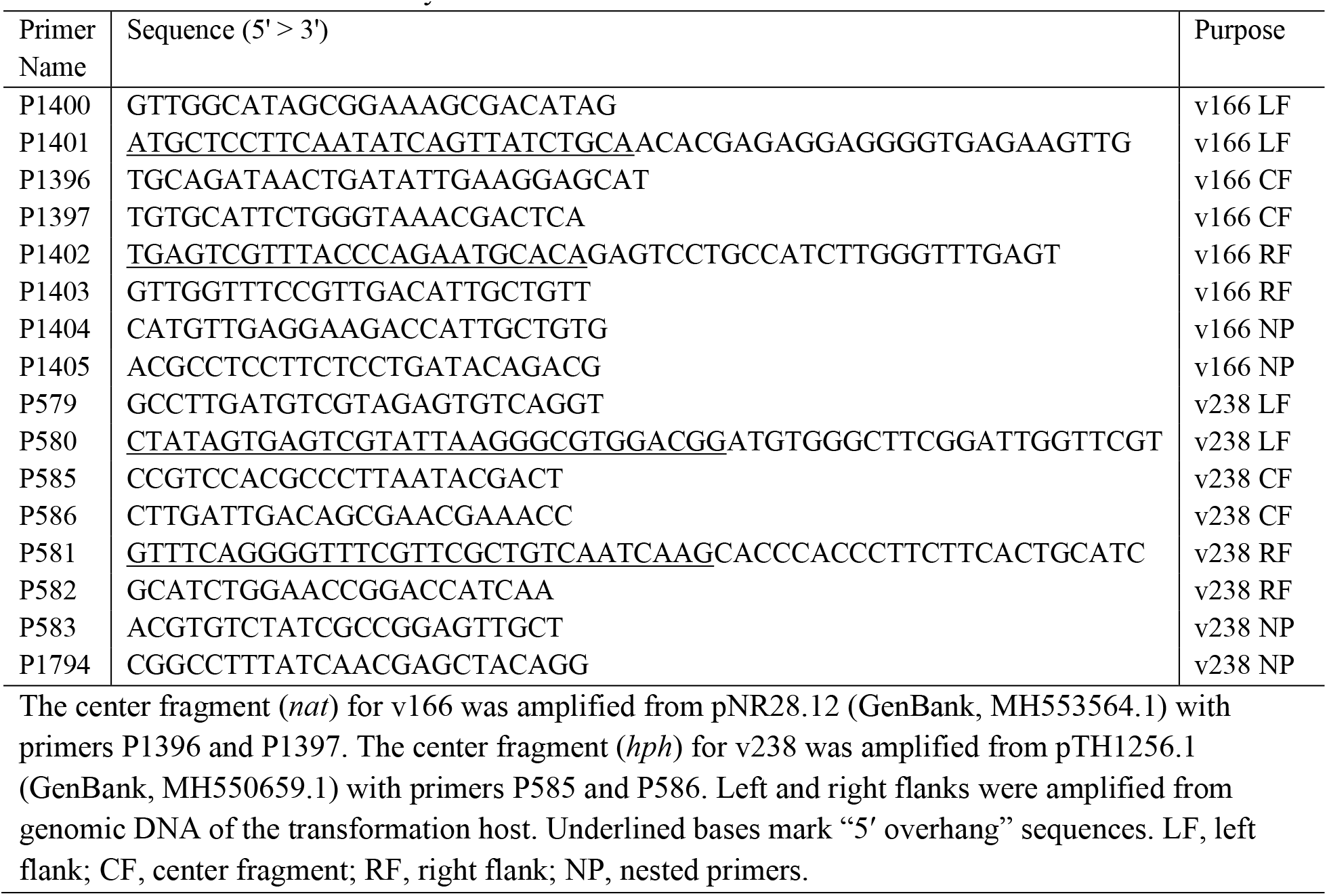
Primers used in this study

### Transformation and homokaryon isolation

Transformation of conidia was adapted from the method described by Margolin et al. (1997). Approximately 1-week old conidia were collected into 30 ml of ice-cold 1 M sorbitol and filtered through a 100 μm cell strainer (Corning 352360). The filtered conidia were pelleted by centrifugation in a swinging bucket rotor at 2000 × g, 4°C. The supernatant was removed and the conidial pellet was resuspended to 1 billion conidia/ml in ice-cold 1 M sorbitol. A 100 μl aliquot of resuspended conidia was then mixed with approximately 300 ng of transformation vector in 10 mM Tris-HCL (pH 8.5). The conidia/DNA mixture (<110 μl) was transferred to a 0.1 cm gap-width electroporation cuvette (BTX model 613) and electroporated at 1500 volts with an Eppendorf Eporator. A 750 μl aliquot of ice-cold 1 M sorbitol was added to the electroporated conidia immediately after the pulse. The electroporated conidia were then transferred to a 4 ml volume of RM in a 50 ml conical tube. This “recovery culture” was incubated at 32°C for 3.5 hours with shaking at 75 rpm, after which 500 μl and 1000 μl aliquots of the recovery suspension were transferred to 10 ml of molten (50°C) TA for plating on 20 ml BA containing nourseothricin sulfate (55 μg/ml) or hygromycin-B (300 μg/ml) in 100 mm culture plates. The transformation cultures were then incubated at room temperature for 12 hours before transfer to a 32°C incubator for an additional incubation period (3 or 4 days). Colonies were then transferred from the transformant plates to 100 × 16 mm culture tubes containing 2 ml of slanted VM with nourseothricin sulfate (45 μg/ml) or hygromycin-B (200 μg/ml). Homokaryons were derived from heterokaryotic *mus-51*^Δ^ transformants by culturing on SC-IAA to generate microconidia for purification with the filtration method of Ebbole and Sachs (1990).

## Results and Discussion

The *mus-51^+^* gene in *N. sitophila* is flanked by genes *ncu08291* and *ncu08289* (Figure 1A), which is similar to situation in *N. crassa* (Galagan *et al.* 2003). We designed a transformation vector, vector 166 (v166), to replace *mus-51* with *nat*. Transformation of FGSC 4746 with v166 produced 47 nourseothricin-resistant transformants. A preliminary screen of the transformants by conidial PCR (Henderson *et al.* 2005) with primers designed to detect replacement of *mus-51* with *nat* identified seven *mus-51*^Δ^ candidates (data not shown). A homokaryotic strain, ISU-4636, was obtained from one of these candidates and PCR was used to confirm that ISU-4636 possesses the *mus-51*^Δ^ genotype (Figure 1B).

**Figure 1.**
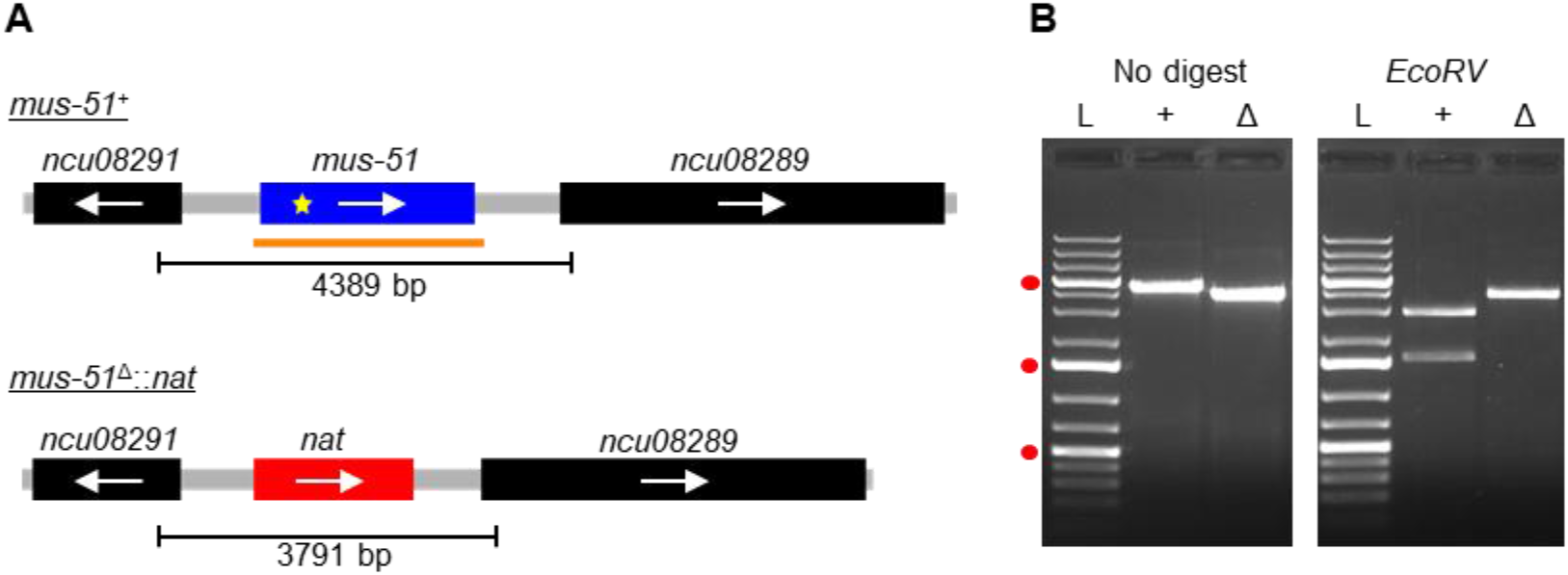
Deletion of *mus-51* from *N. sitophila* FGSC 4746. A) A diagram of the *N. sitophila mus-51^+^* locus is shown above a diagram of *mus-51*^Δ^ locus. Primers P1400 and P1403 amplify a 4389 bp PCR product from the *mus-51*^+^ allele and a 3791 bp PCR product from the *mus-51*^Δ^ allele. The location of a diagnostic *Eco*RV-restriction site within *mus-51^+^* is marked with a yellow star. B) Primers P1400 and P1403 were used to analyze the *mus-51* locus in FGSC 4746 and ISU-4636 by PCR. PCR products were analyzed before *EcoRV*-digestion (left image) and after *EcoRV*-digestion (right image). The results are consistent with *mus-51^+^* and *mus-51*^Δ^ genotypes for FGSC 4746 and ISU-4636, respectively. Lanes: L, GeneRuler 1kb Plus DNA Ladder (Thermo Scientific); +, FGSC 4746; Δ, ISU-4636. Red circles mark positions of 5000 bp, 1500 bp, and 500 bp size markers (top to bottom).

To determine if *mus-51* deletion increases TTI efficiency in *N. sitophila*, we used a cyclosporin (CSA)-resistance assay (Smith *et al.* 2016). The CSA-resistance assay involves transformation of *N. sitophila* strains with a vector designed to replace the *csr-1* gene with *hph*. Previous research in *N. crassa* has shown that strains with a functional *csr-1* allele are sensitive to CSA, while those with a null allele are resistant (Bardiya and Shiu, 2007). Predicting that the relationship between *csr-1* and CSA would exist in *N. sitophila*, we transformed FGSC 4746 (*mus-51^+^*) and ISU-4636 (*mus-51*^Δ^) with vector 238 (v238; designed to replace *csr-1* with *hph*) and isolated fifty hygromycin-resistant transformants from each transformation host. The transformants were then screened for resistance to CSA on CSA-containing medium. While only 9 of 50 transformants from FGSC 4746 demonstrated significant growth on CSA-containing media (Figure 2A), nearly all (49 of 50) of the ISU-4636-derived transformants demonstrated resistance to the compound (Figure 2B).

**Figure 2.**
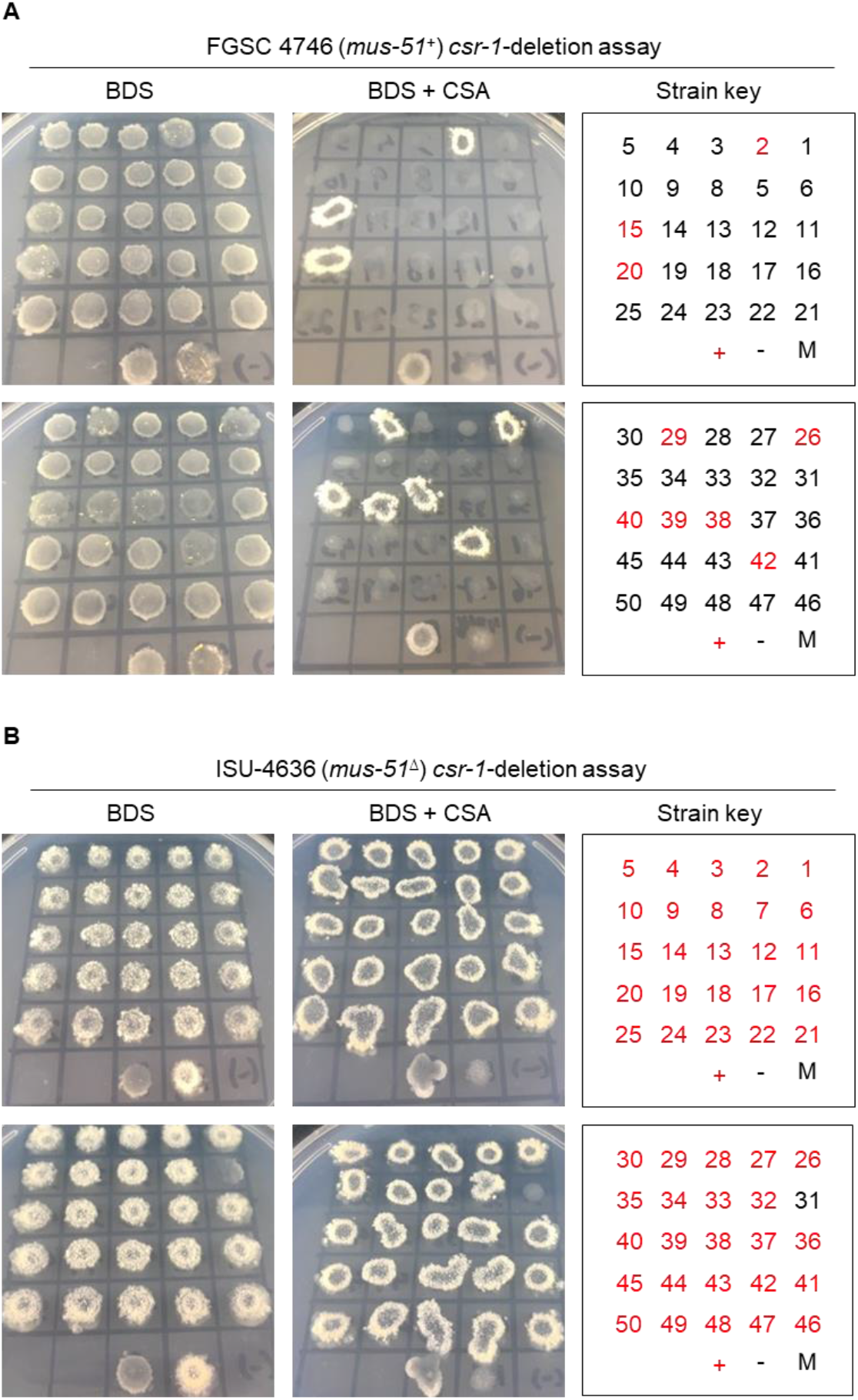
*N. sitophila* TTI efficiency increases fivefold after deletion of *mus-51^+^*. A) FGSC 4746, an *N. sitophila mus-51*^+^ strain, was transformed with vector 238, a vector designed to replace *csr-1*^+^ with *hph*. Fifty hygromycin-resistant transformants were isolated and examined in a cyclosporin A (CSA)-resistance assay. Conidial suspensions (3 μl) of each transformant were transferred to culture plates containing BDS with or without CSA. Images of the culture plates were taken two days after incubation at 32°C. Strain designations: +, P8-65; -, FGSC 4746; 1-50, transformants. Red font is used for strains that were scored as resistant to CSA. B) Similar to panel A, except that ISU-4636 was used as the transformation host. Strain designations are similar to those used in panel A except that “-” refers to ISU-4636.

To confirm that the CSA-resistant phenotype detected with the media-based screen was caused by replacement of *csr-1* with *hph*, the *csr-1* locus was analyzed by PCR in eight transformants from each transformation host. With respect to the FGSC 4746-dervied transformants, we examined four CSA-sensitive and four CSA-resistant transformants and found a perfect correlation between CSA-resistance and the presence of *csr-1*^Δ^ (Figure 3, A and B). With respect to the ISU-4636 transformants, we only isolated a single putative CSA-sensitive transformant (#31), therefore we examined this transformant along with seven CSA-resistant transformants. Interestingly, all eight of the transformants displayed the *csr-1*^Δ^ genotype, even the supposedly CSA-susceptible transformant (#31). It seems that the growth pattern of transformant 31 in the media-based CSA-screen resulted in a “false negative”. Although we have not investigated transformant 31 further, it may possess a growth defect that contributed to the false negative result (note the atypical growth pattern of transformant 31 on BDS without CSA, Figure 2B).

**Figure 3.**
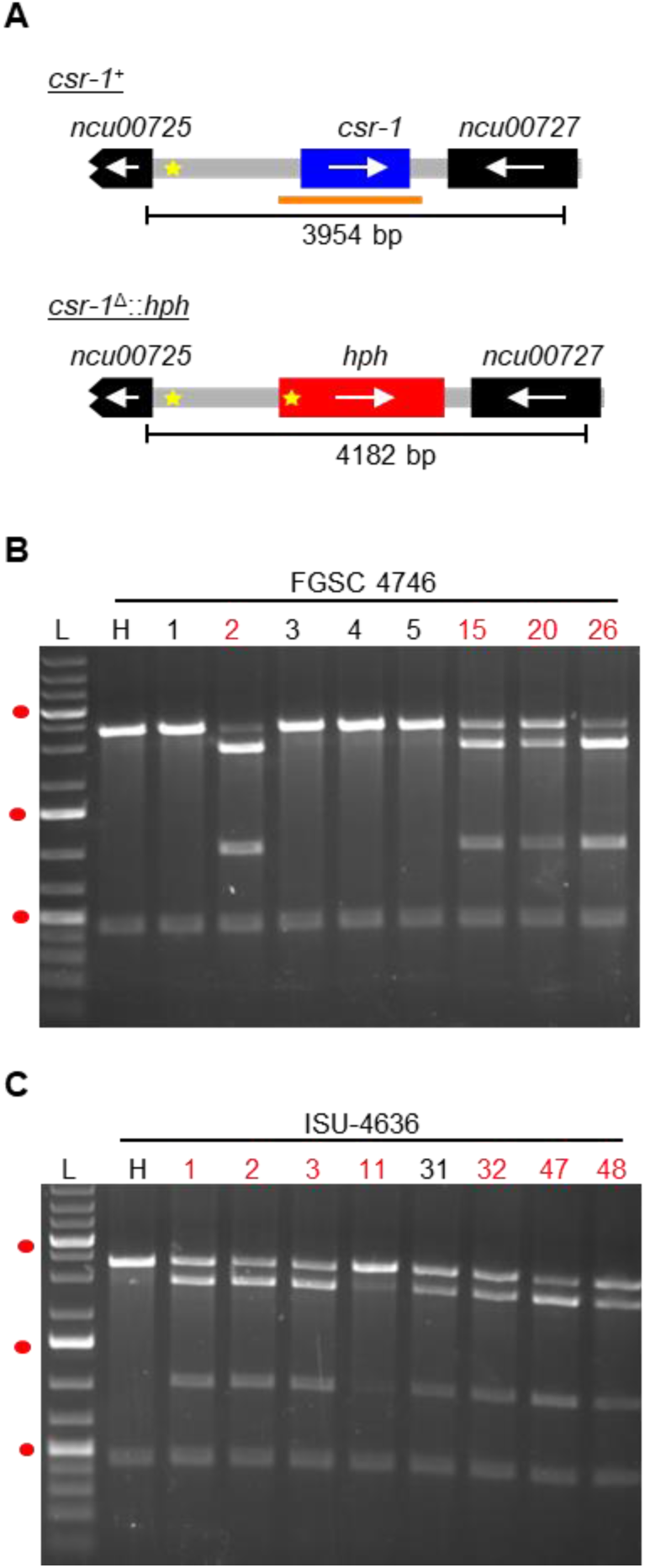
Loss of *mus-51*^Δ^ correlates with increased TTI efficiency in *N. sitophila*. A) A diagram of the *csr-1^+^* locus is shown above a diagram of the *csr-1*^Δ^ locus. *Eco*RI-restriction endonuclease recognition sites are marked with yellow stars. Primers P579 and P582 amplify a 3954 bp PCR product from the *csr-1*^+^ allele and a 4182 bp PCR product from the *csr-1*^Δ^ allele. B) Primers P579 and P582 were used to analyze the *csr-1* locus in FGSC 4746 (H) and eight hygromycin-resistant transformants (1, 2, 3, 4, 5, 15, 20, and 26) obtained by transformation of FGSC 4746 with v238. PCR products were digested with *Eco*RI before analysis by gel electrophoresis. An image of the ethidium bromide-stained gel is shown. L, GeneRuler 1kb Plus DNA Ladder (Thermo Scientific). Red circles mark positions of 5000 bp, 1500 bp, and 500 bp size markers. Red font is used for lane numbers of transformants that were found to be resistant to cyclosporin A (see Fig. 2). C) Primers P579 and P582 were used to analyze the *csr-1* locus in ISU-4636 (H) and eight hygromycin-resistant transformants (1, 2, 3, 11, 31, 32, 47, and 48). PCR products were digested with *Eco*RI before analysis by gel electrophoresis. Labeling is similar to that used in Panel B. Note that the transformants in panels B and C are likely to be heterokaryons (transformed and untransformed nuclei in each transformant).

The results reported above indicate that deletion of *mus-51* increases TTI-efficiency in *N. sitophila*. FGSC 4746, which is the parent strain of ISU-4636, is an *N. sitophila* isolate from Tahiti. To facilitate research on *N. sitophila* strains from different populations, we constructed a *mus-51*^Δ^ strain called ISU-4637 with methods nearly identical to those used to construct strain ISU-4636. The one difference is that we used W1426 (Jacobsen *et al.* 2006) as the transformation host (instead of FGSC 4746). *N. sitophila* W1426 was originally isolated in Italy and will be described in more detail in a future work (H. Johannesson, personal communication). While we have used ISU-4637 in transformation experiments, we have not measured TTI-efficiency, although we have no reason to suspect that ISU-4637 is inferior to ISU-4636 for use in experiments requiring the integration of a transgene into a specific location of the *N. sitophila* genome.

## Conclusion

In summary, our results indicate that the deletion of *mus-51* increases TTI-efficiency in *N. sitophila*. Strains ISU-4636 and ISU-4637 can be obtained from the Fungal Genetics Stock Center as FGSC ##### and FGSC #####, respectively.

## Acknowledgements

We thank Hanna Johannesson, Jesper Svedberg, and Aaron Vogan for *N. sitophila* strains (FGSC 4746 and W1426) and sequences of the *mus-51* and *csr-1* loci. This project was supported by a grant from the National Science Foundation (MCB# 1615626).

